# Polynomial Trajectory Compression for Protein Language Model Embeddings

**DOI:** 10.64898/2026.06.05.730461

**Authors:** Harshita Sahni, Xin Chen, Trilce Estrada

## Abstract

Protein language models (PLMs) generate rich, layer-wise embeddings that capture diverse biological information but are expensive in terms of storage and computation at scale. In this work, we propose a compact surrogate representation for PLM embeddings across transformer layers using low-dimensional PCA projections and cubic polynomial trajectories. This approach enables efficient storage and on-demand reconstruction of these protein-level embeddings at any layer without rerunning the PLM. We evaluate our method on two downstream tasks: protein-protein interaction and subcellular localization using ESM-35M and ESM-3B PLM. We show that the surrogate embeddings achieve high reconstruction fidelity while reducing storage and computational requirements significantly. The new approach also retains downstream task prediction performance compared to original embeddings. Our approach provides a scalable and practical solution for large-scale protein embedding storage and reuse.

## 1 Introduction

Protein language models (PLMs) have emerged as a powerful tool for learning representations directly from long protein sequences. They are architecturally similar to general purpose language models. However, PLMs use transformer architectures that are trained with self-supervised techniques such as masked language modeling on large protein sequence datasets. As protein sequences are processed through successive transformer layers, the model generates intermediate hidden representations, or embeddings, that capture increasingly rich biological information. By training on millions to billions of sequences, PLMs learn contextual representations that capture a wide range of biological properties without requiring explicit structural or functional annotations. The embeddings produced by PLMs encode meaningful information related to protein sequence, structure and function. They are widely used across downstream biological tasks such as secondary struc-ture and contact prediction, protein-protein interaction (PPI) prediction, subcellular localization, enzyme classification, and variant effect prediction [1]. As a result, PLM embeddings have increasingly replaced features such as position-specific scoring matrices and physicochemical descriptors, serving as input for many protein analysis pipelines.

Transformer-based PLMs generate a sequence of layer-wise embeddings. Different layers capture different aspects of the protein. Early layers tend to better capture the local sequence pattern and deeper layers are better at encoding structural motifs. Downstream tasks benefit from embeddings extracted at different layers or their combinations. This has motivated many approaches to extract and store embeddings from multiple layers. However, when applied at scale, i.e. for large protein datasets, this approach introduces significant practical challenges. Storing layer-wise embeddings incurs substantial storage costs. The total storage cost associated with the protein dataset grows proportionally with the length of each protein, the number of layers in the PLM model, the total number of protein samples, and the dimensionality of the output embedding. However, if we do not store the layer embeddings, extracting these layer-wise embeddings for each protein requires computationally expensive forward passes through deep transformer models, often demanding large scale GPU resources. These costs are further amplified in large scale downstream tasks, such as PPI prediction, where embeddings may be computed repeatedly. For example, a model like ESM2-35M [2] with 13 layers and 480-dimensional embeddings, storing all layer-wise representations, requires 6,240 floating-point values per protein (24.4 KB). Larger models such as ESM2-3B [2], with 36 layers and 2,560-dimensional embeddings, demand 92,160 values per protein (368.6 KB) in addition to the transformer size. For a large dataset that contains *∼*1 million protein sequences, gigabytes or terabytes of storage will be required. This becomes a bottleneck as many biological tasks require precomputed embeddings that are reused across many models and various downstream tasks. A single forward pass for larger models like ESM-3B takes *≈* 87.4 milliseconds on a GPU. At a scale of 1 million proteins, common in large proteome studies, it can take about 24 hours per experimental run. Moreover, the same protein embeddings from different PLM layers can be used for different downstream tasks such as PPI prediction, function annotation, and many others. Therefore, recomputing the same embeddings for different experimental runs or tasks can waste significant amounts of resources.

In this work we address this challenge by introducing a technique that compresses embeddings across layers for efficient storage. Instead of storing embeddings independently at each layer, we model the entire trajectory with compact polynomial functions and reconstruct any layer on demand. Our contributions are as follows:

1. We propose a novel compression framework that models the layer-wise trajectory of protein-level (mean-pooled) PLM embeddings using compact polynomial functions, enabling any layer’s representation to be reconstructed on demand without storing all intermediate layers.
2. Our approach substantially reduces the storage and computational burden of full layer-wise protein-level (mean-pooled) PLM embeddings, making large PLMs more accessible to researchers without extensive infrastructure, while still producing embeddings suitable for downstream tasks.
3. We validate our surrogates across two downstream tasks: protein-protein interaction prediction and subcellular localization; and show that surrogate embeddings retain task performance.

The paper is organized as follows: Section 2 provides a literature review. Section 3 introduces our compression approach and details the methodology used in this work. Section 4 discusses the results and demonstrate the effectiveness of our compression approach on two downstream tasks. Finally, Section 5 concludes the paper and outlines directions for future work.

## 2 Related Work

Being at the intersection of protein language models (PLMs), representation analysis, and embedding compression, below we present a literature review across these domains respectively.

### Protein Language Models

Protein language models (PLMs) use transformer architectures on protein sequences. PLMs are typically trained using self-supervised techniques like masked language modeling that are similar to models like BERT [3]. These models learn to predict masked amino acids from the unmasked context, trained on millions to billions of protein sequences from UNIPROT [4]. They learn powerful representations for biological sequences that capture functional and structural information. These representations or embeddings can be transferred to many downstream biological tasks, replacing handcrafted features such as position-specific scoring matrices or physicochemical descriptors. Several PLM architectures have been proposed in recent years and have been used for downstream tasks as listed above [1] [5]. Early models such as UniRep demonstrated that unsupervised sequence models can capture meaningful biochemical properties [6]. More recently, transformer-based models like ProtBERT [7], Prottrans [8], and the ESM (Evolutionary Scale Modeling) family [2] [9]. comprise tens of millions to billions of parameters [10] [11]. Increasing model size has consistently been associated with improved representation quality and downstream task performance. Work from Rives et al. systematically studied the effect of scaling transformer models, and showed that important biological properties are captured better when model size and depth increase [9]. Their work also demonstrates that PLMs implicitly encode secondary structure, residue–residue contacts, and functional annotations through different layers. Additionally, Elnaggar et al. [8], and Rui et al. [12] showed that transformer-based PLMs consistently outperform traditional handcrafted features across diverse downstream tasks, including structure prediction and PPI classification.

### Layer-wise Representation Analysis

Several studies have examined how representations extracted from different layers of protein language models behave across different down-stream tasks. Vig et al. [13] showed that attention heads in protein transformers align with meaningful biological signals, such as residue contacts and conserved motifs. In general, earlier layers capture local sequence patterns, while deeper layers encode longer-range and more abstract relationships, suggesting a hierarchical organization of information across network depth. Similarly, Rao et al. [14] introduced the TAPE benchmark and showed that embeddings from intermediate layers often outperform final-layer embeddings for structure-related tasks, such as secondary structure and contact prediction. More recent analyses have compared embeddings across layers and models. Kumar et al. [15] systematically evaluated representations from different layers of multiple protein language models for kinase classification and found that no single layer is universally optimal across tasks. They conclude that mid-to-late transformer layers performed better than the last layer. These findings further support the idea that biological information is distributed across layers rather than best represented through the final layer. Storing embeddings from all the layers can be expensive. In our work, we provide a compression technique that allows storing embeddings from all layers at low cost. This will enable on-demand and efficient embedding extraction of any layer.

### Model Compression and Distillation

The growing size of PLMs has motivated research into reducing their computational and storage costs. Knowledge distillation, originally introduced by Bucilua et al. [16] and later generalized by Hinton et al. [17], transfers knowledge from a large teacher model to a smaller student model and has been widely used to compress deep neural networks. For protein models, dimensionality reduction blocks that efficiently compress the latent space into a more parsimonious intermediate-level representation are widely popular [18]. Other methods focus on reducing redundancy in model parameters or activations. Low-rank approximations and factorization-based methods have been proposed to compress transformer models by exploiting the low-dimensional structure of hidden representations [19] [20]. However, these approaches typically require fine-tuning and still rely on full forward passes through the model during inference.

In contrast to model-level compression, our work operates directly on extracted layer-wise embeddings. After a single forward pass to obtain embeddings for a protein, we exploit structure in the representation space itself to construct compact surrogate representations. This enables efficient storage and reconstruction of layer-wise embeddings without modifying model parameters or requiring repeated inference through the language model.

### Representation Geometry and Smoothness

Beyond task performance, several studies examine the geometric structures of neural representations across layers in deep models. Raghu et al. introduced the SVCCA architecture to analyze deep representations across layers, and concluded that lower layers learn first [21] and proposed a novel technique to compress and speed up neural training. Kornblith et al. further proposed similarity metrics for comparing representations and demonstrated substantial redundancy and alignment across layers in deep networks [22].

However, in the context of protein language models, the geometric structure of layer wise embeddings remains unexplored. Understanding how protein representations evolve across layers and whether this evolution follows smooth, predictable trajectories, and if embeddings of all layers can be compressed using surrogate models lays the foundation of this work.

## 3 Method

Through this work we investigate whether embeddings generated through a PLM can be represented using a compact surrogate representation to reduce storage requirements while preserving biologically relevant information. Figure 1 illustrates our workflow. In the following sections, we describe the embedding extraction pipeline and surrogate modeling approach used to approximate layer wise embeddings.

**Figure 1:**
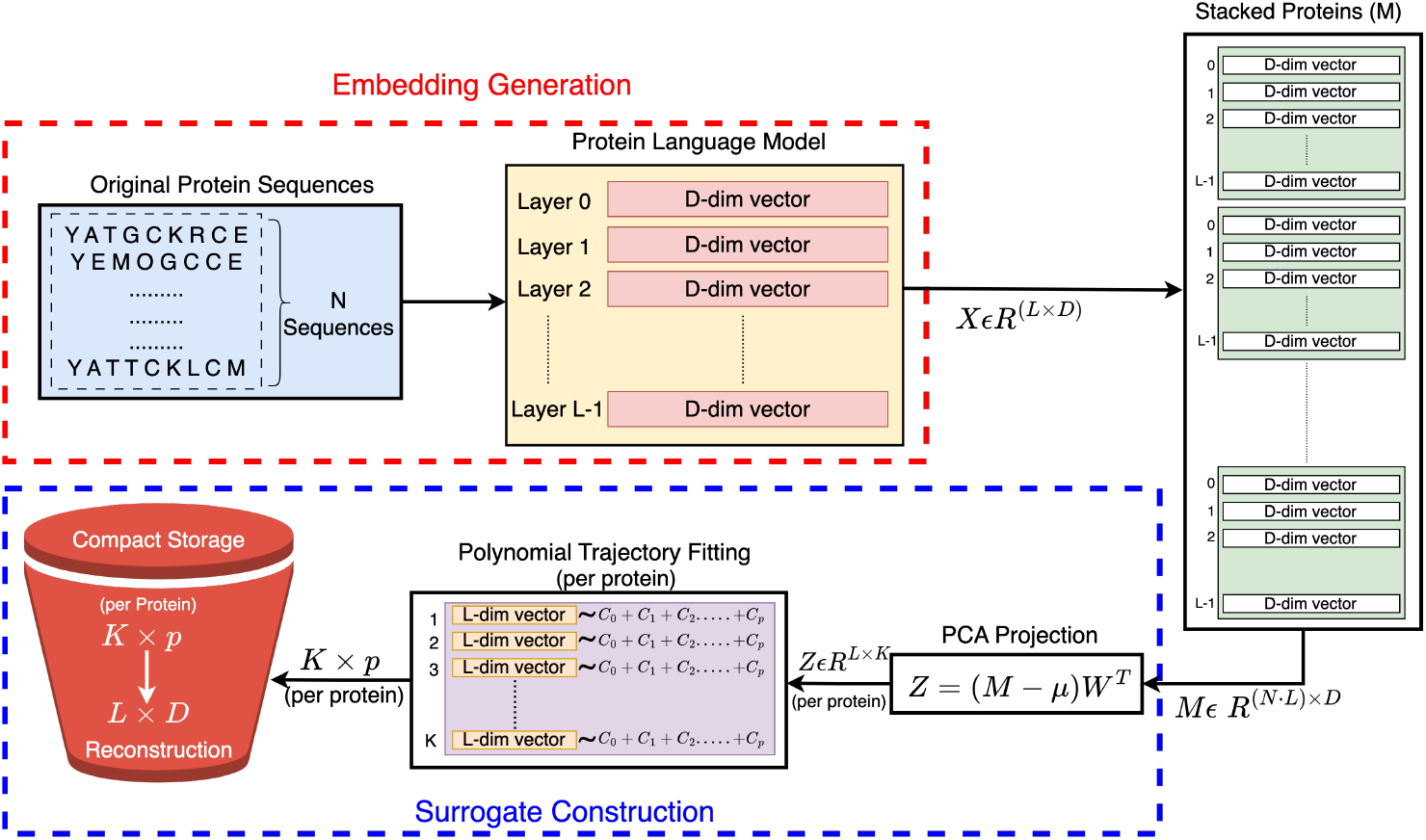
Surrogate Compression of PLM Embeddings.

### 3.1 Embedding Generation

For each protein sequence in our dataset, we perform a forward pass and extract layer wise embeddings from two protein language models (i.e., ESM2-35M and ESM2-3B) in the ESM2 family ^1^. ESM2-35M, has 12 transformer layers and 480-dimensional embeddings. It serves as our primary model for detailed analysis in this work. ESM2-3B has 36 transformer layers and 2560-dimensional embeddings, is used to demonstrate scaling efficiency of our approach to larger architectures. Both models were pretrained on approximately 65 million protein sequences from UniRef50 [23], making them representative of state-of-the-art PLMs widely used in computational biology.

ESM2-35M produces embeddings at layers 0 through 12 (13 layers total and *≈*35 million parameters), while ESM2-3B produces embeddings at layers 0 through 36 (37 layers total and *≈*3 billion parameters). Layer 0 corresponds to the input embedding layer, and subsequent layers represent progressive transformations through the transformer blocks. At each transformer layer, the model outputs per-residue representations. For a protein sequence of length *Y*, the output at layer *ℓ* is a matrix of shape (*Y, D*), where *D* denotes the embedding dimension (480 for ESM2-35M and 2560 for ESM2-3B). To obtain a fixed-length representation for each protein, we apply mean pooling across the sequence dimension at every layer. Special tokens, including beginning-of-sequence, end-of-sequence, and padding tokens, are excluded prior to pooling. The protein-level (mean-pooled) representation at layer *ℓ* is computed as: 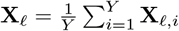 where **X**_*ℓ,i*_ represents the embedding of residue *i* at layer *ℓ*. This produces a single *D*-dimensional embedding vector per protein per layer. Stacking embeddings across all layers yields a layer-wise representation matrix **X***∈* ℝ^*L×D*^ for each protein, where *L* is the total number of transformer layers.

For protein sequences exceeding the model context limit of 1024 residues, we use a sliding-window approach. Sequences are partitioned into overlapping windows of 1024 residues with a stride of 512, with the final window adjusted to ensure coverage of the sequence end. Each window is processed independently to extract layer-wise embeddings, and the final protein representation is computed as a length-weighted average across all windows. This ensures that all residues contribute proportionally to the final embedding regardless of sequence length.

Protein embeddings are computed using the ESM models through their PyTorch implementation. We used single-precision (FP32) computation on NVIDIA L40S GPUs to reduce memory requirements. To avoid GPU out-of-memory errors we used a batch size of 4 for ESM2-35M. For ESM2-3B, which requires approximately 11GB of GPU memory, we used a batch size of 2. To parallelize embedding extraction, we distributed proteins across multiple GPU shards using a deterministic round-robin assignment and processed each shard independently. Embeddings were processed in chunks of 200 proteins and stored in compressed NumPy archives (.npz format) with protein’s *UniProt-ID* as keys and layer-wise embeddings as values. For ESM2-35M, we needed to store 6,240 floating-point values per protein (13 layers *×* 480 dimensions), totalling 25 GB for 1M proteins at FP32; for ESM2-3B, it was 92,160 floating-point values per protein (36 layers *×* 2560 dimensions) requiring 369 GB for the same dataset. These storage requirements motivate our compression approach described in Section 3.2.3.

### 3.2 Surrogate Construction

Our surrogate construction approach transforms high-dimensional layer wise embeddings into a low dimensional latent space that enables reconstruction at any layer. The method consists of three steps: (1) dimensionality reduction via principal component analysis (PCA) to identify a lower rank subspace spanned by the top *K* principal components, (2) *p™*1 dimensional polynomial fitting to model smooth trajectories through this subspace, and (3) storing compact, per protein, coefficients that enable on-demand reconstruction. This approach reduces storage requirements from *L ×D* floating-point values per protein to *K×p* coefficients, where *K << D* and *p* is defined by the coefficients in a polynomial. The choice of (*K, p*) is task-dependent and selected on the validation set: for PPI prediction we use *K* = 64, *p* = 4 (cubic) across both ESM models; for the subcellular localization task we use *K* = 128, *p* = 4 for both ESM-35M and ESM-3B, (see Section 4.3).

#### 3.2.1 PCA Projection

To identify the low-dimensional subspace, we apply PCA to the generated embeddings for the proteins in the training set processed through ESM models. It is well-suited for our task because it identifies the directions of maximum variance in the data, ensuring that the low-dimensional representation captures the most significant variations in how embeddings change across layers. It also offers an efficient linear projection that can be applied to new proteins at test time without retraining. The method proceeds as follows.

Given a model with *L* layers and *D*-dimensional embeddings, and a training dataset with *N* proteins, for which 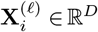 is the embedding of protein *i* at layer *ℓ*. We construct a data matrix **M** *∈* ℝ (*N ·L*)^*×D*^ by stacking the layer-wise embeddings from all *N* training proteins. We then compute the mean embedding of the dataset as 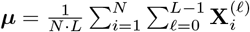 and the covariance matrix 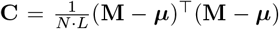. PCA identifies the top *K* eigenvectors of **C**, which we organize as rows of the projection matrix **W** *∈* ℝ^*K×D*^.

For each protein *i* with embedding 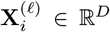 at layer *ℓ*, we project this original layer-wise embedding onto the *K*-dimensional PCA subspace:

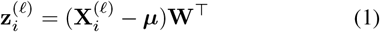

where 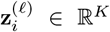 is the PCA projection for protein *i* and layer *ℓ*. Stacking across all layers and all proteins **Z** *∈* ℝ^(*N·L*)*×K*^ contains the PCA-projected coordinates for all proteins. We fit PCA using only training embeddings yielding (***µ*** and **W**). For validation and test proteins we apply the same transformation to their layer-wise embeddings as described in equation (1). This ensures that no information from validation or test sets influences the choice of low-dimensional subspace, preventing data leakage. Hence, any new protein with embedding 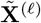 and dimension *L × D* yields a PCA projection 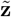 of size: *L × K* in accordance with equation (1).

#### 3.2.2 Polynomial Trajectory Fitting

Once we have projected per-protein embeddings for each layer into a low-dimensional PCA space, we observe that a protein’s trajectory across layers follows a continuous path (Appendix Figure 4). This occurs as each transformer layer adds to its input representation rather than replacing it, so the representation changes gradually across layers. This smoothness (few abrupt changes) motivates our use of polynomial functions to model these trajectories. We normalize layer indices between [*−*1, 1] by mapping the initial layer to *t*_0_ = −1 and the final layer to *t*_*L−*1_ = 1. Our process is formulated as follows:

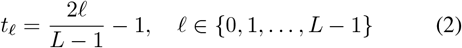

where *ℓ* is the layer index starting from zero and *L* is the total number of layers. This ensures numerical stability during polynomial fitting and keeps all the values within a bounded range.

After PCA projection, each protein *i* is represented by a matrix **z**_*i*_ *∈* ℝ^*L×K*^, where each row 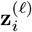 contains the coor-dinates of the top *K* principal components at layer *ℓ*. For each principal component *k ∈* {1, …, *K*}, we obtain a sequence of values across layers:[*z*_0,*k*_, *z*_1,*k*_, …, *z*_*L−*1,*k*_], which describes how the coordinate associated with component *k* evolves throughout the transformer layers. For example, in ESM2-35M with *L* = 13, this corresponds to 13 observed values (one per layer) for each principal component.

To model this layer-wise trajectory, we fit a polynomial function for each protein and principal component. Specifically, the polynomial maps the normalized layer position *t*_*ℓ*_ to the predicted PCA coordinate value:

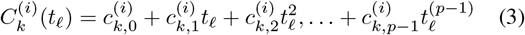

where 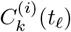 predicts the PCA coordinate for dimension *k* of protein *i*, where *k ∈* 1, …, *K* at normalized layer position *t*_*ℓ*_, and **c**_*k,j*_ is the *j*-th polynomial coefficient for the PCA dimension *k* of protein *i*, for *j ∈*{0, 1, …, *p−*1}. These polynomial coefficients are learned using ordinary least squares [24] on the observed PCA coordinates *z*_0,*k*_, *z*_1,*k*_,…, *z*_*L−*1,*k*_ across all *L* layers to minimize the reconstruction error. Therefore, for each protein represented through *L* layers and each layer with *K* PCA components can now be represented by *K× p* coefficients (values of *K* used in this work range from 64 to 128 and values of *p* = 4).

#### 3.2.3 Surrogate Reconstruction

The surrogate representation for each protein consists of *K* sets of *p* polynomial coefficients, totaling *K×p* floating-point values. For models with *K* = 64 and *p* = 4, this requires 256 floating-point values per protein (1 KB in FP32).

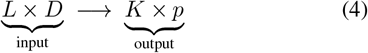

Reconstruction of any embedding from the surrogate at any layer *ℓ* for any protein *i* can be done through the following steps:

##### Step 1: Normalize layer index

**Input:** Target layer *ℓ ∈* {0, 1, …, *L* − 1}

**Output:** Scalar *t*_*ℓ*_ *∈* [−1, 1], computed in Equation (2).

##### Step 2: Evaluate *K* polynomials at *t*_*ℓ*_

**Input:** Scalar *t*_*ℓ*_ and stored coefficients 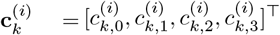 for each *k ∈* {1, …, *K*}

**Output:** *K* scalar values 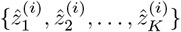, computed as:

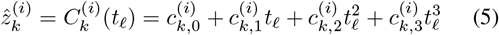

##### Step 3: Assemble reconstructed PCA vector of size *K*

**Input:** *K* scalar values 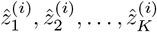 from Step 2

**Output:** Reconstructed PCA coordinate vector of *K* dimensions at layer 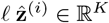 :

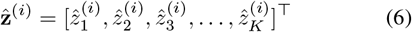

##### Step 4: Project to original embedding space

**Input:** 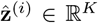 from Step 3, global PCA components **W** *∈* ℝ^*K×D*^, and PCA mean ***µ*** *∈* ℝ^*D*^

**Output:** Reconstructed embedding 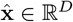 at layer *ℓ*:

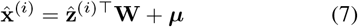

## 4 Evaluation

In this section we will present and discuss our results. We evaluate our surrogate compression technique in three ways: (1) reconstruction quality of compressed surrogate embeddings; (2) effectiveness of compressed embeddings on a PPI prediction and subcellular localization downstream tasks; (3) computational and storage efficiency of compressed embeddings during inference. In the following sections we present results for ESM-35M. We also present results for ESM3B that shows that our compression approach can be scaled to larger models.

### 4.1 Datasets

#### PPI prediction

We first apply our approach to a large-scale PPI dataset collected from the study in [25]. This dataset provides experimentally validated interacting and non-interacting human protein pairs and has been constructed to eliminate data leakage between training, validation, and test splits. The final dataset consists of 163,192 training pairs, 59,260 validation pairs, and 52,048 test pairs. The total number of positive pairs is equal to the total number of negative pairs. This setup enables a robust assessment of generalization performance for downstream PPI predictions and allows us to evaluate whether surrogate embeddings preserve biologically relevant interaction signals under strict data leakage constraints.

#### Subcellular localization

For the localization task, we used the DeepLoc 1.0 benchmark [26]. This is a multi-class classification task. Each protein is annotated with one of ten subcellular compartments: cytoplasm, nucleus, extracellular, mitochondrion, cell membrane, endoplasmic reticulum, plastid, Golgi apparatus, lysosome/vacuole, and peroxisome. The dataset is curated from UniProt with experimentally verified localization annotations and is redundancy reduced at 30% sequence identity to limit homology between train and test splits. It contains 11,231 training and 2,773 test proteins drawn from diverse eukaryotic organisms; we carve a stratified 10% validation subset from the training set for hyperparameter selection.

### 4.2 Results and Discussion

First, we quantify how well the surrogate approximation preserves the original embedding using the test dataset. We evaluated this using Cosine Similarity [27] that measures how similar the surrogate embedding is to the original embedding. As shown in Figure 2 (a) most proteins reconstruct well and achieve high cosine similarity values with a mean at 0.9154 and median centered at 0.9273. For ESM-3B (larger transformer model), our cosine similarity scores was even better and ranged from 0.98-0.99. For subcellular localization, where we use a richer surrogate (*K* = 128), reconstruction quality is consistently high across both models: mean=0.98 for ESM-35M and ESM-3B and ranged from 0.92-0.99. We next assess the question “if this level of reconstruction quality is sufficient to perform downstream classification task of PPI and subcellular localization prediction?”

**Figure 2:**
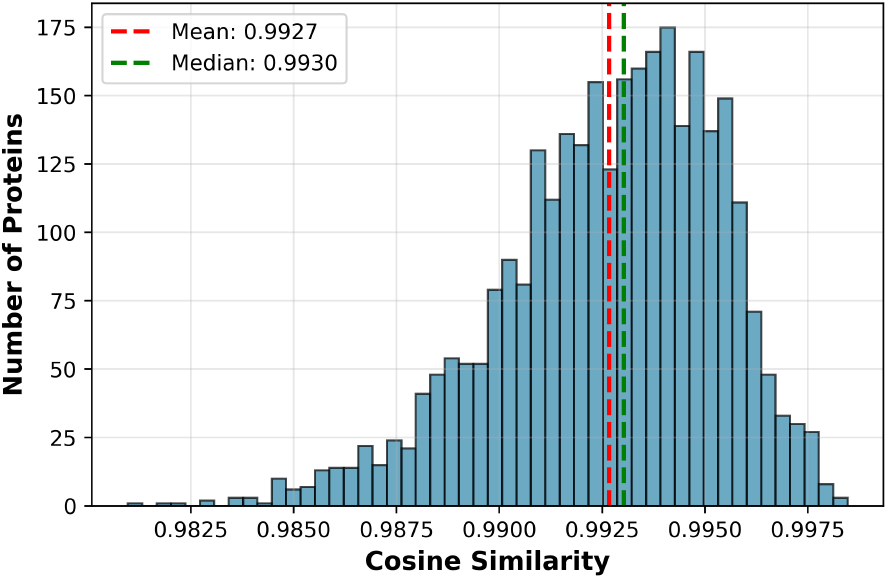
Cosine similarity reconstruction quality: distribution for ESM-3B.

### 4.3 Downstream task evaluation

To evaluate whether the surrogate embeddings preserve biologically relevant information beyond reconstruction quality, we assess their performance on two downstream tasks: protein-protein interaction (PPI) prediction and subcellular localization prediction. PPI is a supervised binary classification task. Given two proteins, Prot1 and Prot2, with their respective sequences, the goal is to predict if the two proteins interact (label 1) or do not interact (label 0). On the other hand, subcellular localization prediction is a single-protein, ten-class classification task. Given a protein with its sequence, the objective is to predict the subcellular compartment in which the protein resides (one of ten classes). Next, we assess whether our surrogate embeddings preserve the biological information and can achieve similar performance on the two downstream tasks. To achieve this, we followed the steps listed below:

- Collect the original embeddings for all layers for validation and test data using ESM models.
- Identify the optimal layer for the specific downstream task using validation data.
- Reconstruct surrogate embeddings for that layer using our method described in section 3.2.3
- Train two separate classifiers, one for the original embeddings and another for the surrogate embeddings, using the layer embeddings selected in the above step.
- Compare prediction performance of the original vs. surrogate embeddings on the test dataset.

#### 4.3.1 Layer Selection for downstream tasks

To identify the most informative layer for the PPI and subcellular localization prediction task, we evaluated each of layers of the PLM. For each layer *ℓ*, we extracted embeddings for all proteins in the training set and trained a logistic regression model, separately for the original and surrogate embeddings. For the downstream tasks, each protein pair is represented by combining the embeddings of the two proteins at layer *ℓ* via the Hadamard product (i.e., element-wise multiplication) [28]. This serves as the input to the logistic regression model, which predicts whether the pair interacts (binary classification). For the subcellular localization task, the layer-*ℓ* embedding of a single protein is used directly as input. The logistic regression model predicts one of ten subcellular compartments (multiclass classification). We then used the validation set to select the hyperparameters: optimal layer, PCA dimension *K*, and polynomial degree *p*. For the PPI prediction downstream task, and based on the validation set, we choose *K* = 64 with a cubic polynomial (*p* = 4) as lower values of K (16, 32) reduce AUROC by up to 0.030 (0.593 vs 0.623), while *K* = 128 offers less than 0.003 improvement (0.620 vs 0.623). Lower-degree polynomials (degree 1, 2) degrade downstream PPI prediction and higher degrees offer no consistent benefit, with AUROC declining to 0.608 at degree 5. This trend was observed across both ESM models. Similarly, the configurations selected on the validation set for subcellular localization are *K* = 128, *p* = 4 for ESM-35M and ESM-3B. These hyperparameters are fixed before evaluation on the test set).

Figure 3 (a) and (b) shows the performance using AUROC, AUPRC, precision, recall and F1 score across different layers. AUROC and AUPRC serve as the primary metrics for ranking and selecting the layers for the PPI prediction task using ESM-35M. As shown in Figure 3, performance gradually increases starting from layer-0 and then plateaus through the middle layers (7-11) and peaks at layer 12. This layer-wise evaluation has an important implication that no single layer is universally optimal for all downstream tasks. This finding further strengthens our approach to store embeddings from multiple layers rather than just the final layer. While layer 12 performs best for downstream task using ESM-35M (based on AUROC score), other tasks may benefit from a different layer or combination of layers. For the subcellular localization task, the layer scan exhibits a similar pattern and we select layer 11 as the best layer.

**Figure 3:**
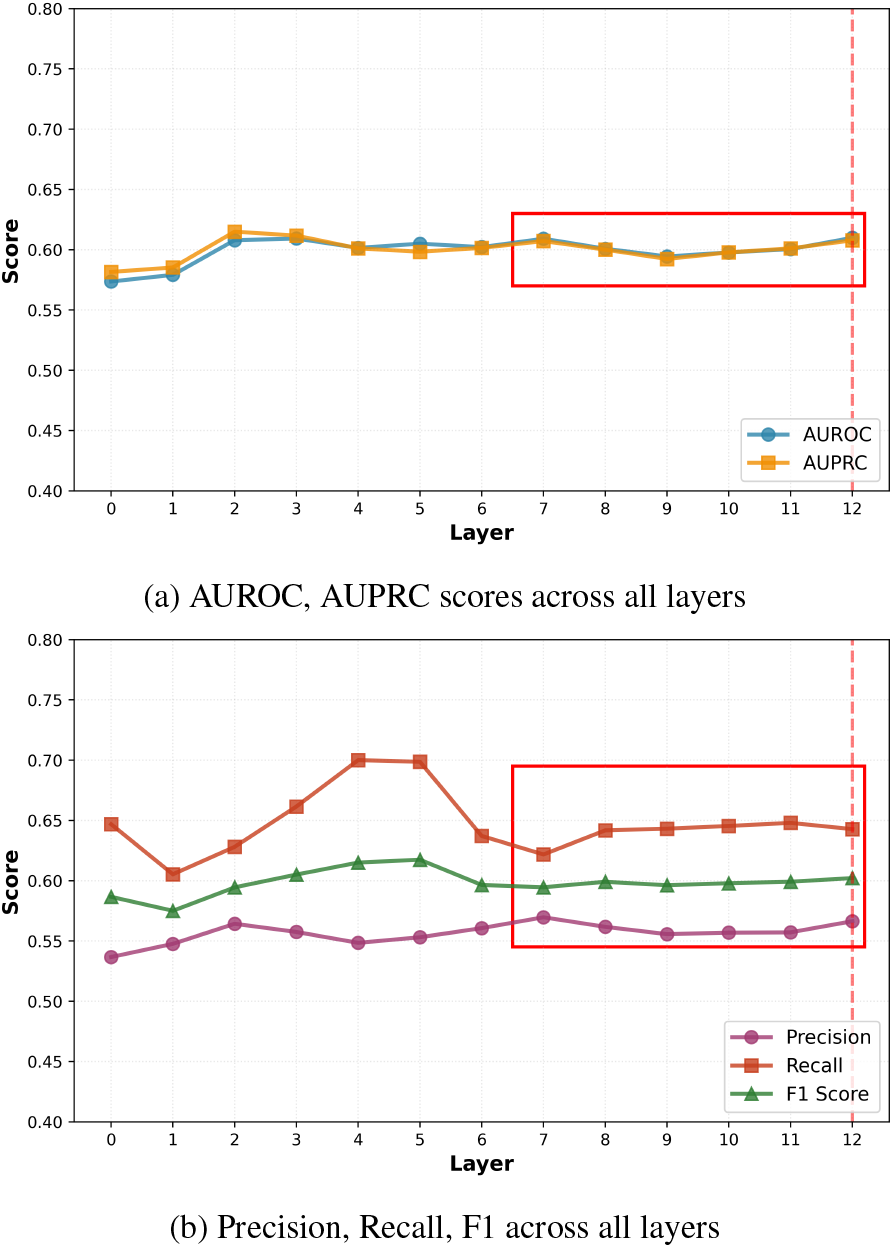
Layer-wise PPI prediction performance using ESM-35M on the validation dataset.

For ESM-3B (total layers = 36), layer 32 works the best for the PPI task and also for localization task (Appendix Figure 6). Although we used a very simple logistic regression model for the PPI prediction task, this study [29] shows that accuracy plateaus at 66% for the same dataset even when complex models such as transformers are used.

#### 4.3.2 Surrogate Embedding Performance

##### PPI prediction

After identifying layer 12 as an optimal layer for the PPI prediction task, we next evaluated whether the surrogate embeddings in this layer preserve the biological information related to the task present in the original embedding. To test this hypothesis, we used the trained logistic regression model on the surrogate embeddings of the training dataset to evaluate its performance on the held-out test dataset. To compare our results with the original embeddings, we used the same training procedure and hyperparameters. Table 1 presents the performance comparison. The surrogate embeddings maintain similar performance and in some cases exceed original embeddings. The surrogate embeddings achieve AUROC of 0.659 compared to 0.648 the original embeddings. These results demonstrate that our polynomial surrogate can work similarly to ESM embeddings despite achieving 96% reduction in data size per protein. To demonstrate the robustness of the task across multiple layers, we also plot the performance metric for layer 7, a middle layer in the transformer (Appendix Figure 5). Importantly, the surrogate embeddings are again comparable to, and slightly higher than the original embeddings with an AUROC of 0.670 compared to 0.653. This robustness across layers provides storage flexibility to the researchers. They can select a subset of layers for any downstream task without committing to store all layers or their original-embeddings. For ESM2-3B AUROC is 0.650 for original embeddings vs 0.647 for surrogate embeddings Table 1). These numbers confirm that our approach is scalable across model sizes.

**Table 1:** Test-set performance of original and surrogate embeddings across two downstream tasks (PPI and Subcellular Localization (SubLoc)) and two PLM scales, with 95% confidence intervals from bootstrapping test predictions (1000 resamples).

| Task | Model | Type | Accuracy | AUROC | AUPRC | F1 |
| --- | --- | --- | --- | --- | --- | --- |
| PPI | ESM-35M (L12) | Original | 0.602 [0.598,0.606] | 0.648 [0.643,0.653] | 0.648 [0.642,0.654] | 0.605 [0.600,0.610] |
|  |  | Surrogate | 0.608 [0.604,0.612] | 0.659 [0.654,0.663] | 0.651 [0.644,0.657] | 0.638 [0.633,0.643] |
|  | ESM-3B (L32) | Original | 0.600 [0.596,0.604] | 0.650 [0.645,0.654] | 0.663 [0.657,0.668] | 0.625 [0.621,0.630] |
|  |  | Surrogate | 0.608 [0.604,0.613] | 0.647 [0.643,0.652] | 0.637 [0.630,0.643] | 0.590 [0.585,0.595] |
| SubLoc | ESM-35M (L11) | Original | 0.749 [0.732,0.766] | 0.934 [0.926,0.943] | 0.794 [0.779,0.812] | 0.738 [0.720,0.755] |
|  |  | Surrogate | 0.726 [0.709,0.743] | 0.925 [0.916,0.934] | 0.769 [0.754,0.788] | 0.710 [0.692,0.728] |
|  | ESM-3B (L32) | Original | 0.817 [0.802,0.830] | 0.957 [0.949,0.964] | 0.856 [0.842,0.873] | 0.812 [0.797,0.826] |
|  |  | Surrogate | 0.818 [0.802,0.832] | 0.954 [0.945,0.961] | 0.851 [0.836,0.867] | 0.810 [0.794,0.825] |

##### Subcellular localization

We followed the same procedure for the subcellular localization task at layer 11 of ESM-35M. Table 1 compares original and surrogate embeddings on the held-out test set. The surrogate retains the original embeddings performance to within *≈*1% AUROC: 0.925 compared to 0.934 for original embeddings. For ESM-3B at layer 32, the surrogate achieves AUROC of 0.954 compared to 0.957 for original, demonstrating minimal degradation in the performance. Across both tasks and scales, surrogate performance stays within *≈*0.011 AUROC of the original.

### 4.4 Computational and Storage Efficiency

Here we will discuss the practical advantages of our approach. We consider two common deployment scenarios: (1) ondemand computation, where embeddings are generated as needed without being previously stored, and (2) pre-computed storage, where embeddings are stored for repeated access.

#### On-demand computation

Table 2 compares inference time for ESM embedding computation Vs surrogate reconstruction. On GPU ESM2-35M requires *≈* 12 milliseconds per protein, while surrogate reconstruction on CPU takes only *≈* 0.0008 milliseconds. Once a protein’s coefficients are computed and stored, every subsequent access reconstructs its embeddings in real-time on a CPU. This can eliminate repeated GPU forward passes across layers, tasks, and experimental runs. Generating coefficients for a new protein still requires a single forward pass. Therefore, the savings come from reuse rather than from the initial encoding. In terms of storage our surrogate model requires only shared PCA mean vector and components, independent of number of proteins. For ESM2-35M (D=480), the surrogate storage is 123KB with *K* = 64 and 246KB with *K* = 128 compared to 320MB for the full PLM. For ESM2-3B (D=2560), it is 650KB for *K* = 64 and 1.3MB for *K* = 128 compared to 26.8GB for the full model.

**Table 2:** Comparison of Inference time per model type.

| MODEL TYPE | DEVICE | RUNTIME (MS) |
| --- | --- | --- |
| ESM2-35M | GPU | $1.220 \times 10^1$ |
| SUR-35M | GPU | $0.026 \times 10^{-3}$ |
| SUR-35M | CPU | $1.200 \times 10^{-3}$ |
| ESM2-3B | GPU | $8.740 \times 10^1$ |
| SUR-3B | GPU | $0.281 \times 10^{-3}$ |
| SUR-3B | CPU | $4.960 \times 10^{-3}$ |

#### Pre-computed storage

Storing all-layer embeddings is the dominant cost at scale. For ESM-35M, full embeddings require 24.4 KB per protein (24.4 GB for 1M proteins), while our surrogate requires only 1.0 KB per protein for PPI (*K* = 64, *p* = 4) and 2.0 KB for subcellular localization (*K* = 128, *p* = 4), yielding 96% and 92% reductions, respectively. The gain is far larger for ESM-3B: full embeddings require 360 KB per protein (360 GB for 1M proteins), while the surrogate requires only 1.0 KB (PPI, *K* = 64, *p* = 4) or 2.0 KB (localization, *K* = 128, *p* = 4), a 99% reduction across both tasks.

For ESM2-35M (*L* = 13, *D* = 480, *K* = 64, *p* = 4): (96% reduction)/protein

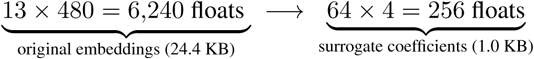

For ESM2-3B (*L* = 36, *D* = 2560, *K* = 64, *p* = 4): (99% reduction)/protein

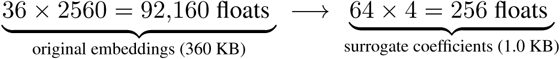

These reductions combine two stages: PCA gives a *D/K* reduction, and polynomial fitting adds a further *≈L/p* factor. PCA-only achieves comparable AUROC (Appendix A.1), so the polynomial step adds extra compression at similar performance.

## 5 Conclusion

In this work we presented a scalable framework for compressing and reconstructing layer-wise PLM embeddings. By modeling embedding trajectories with low-dimensional PCA projections and polynomial surrogates, our approach achieves a significant reduction in storage: 96% for ESM-35M and 99% for ESM-3B models per protein while maintaining high cosine similarities of 0.91-0.99, without rerunning the PLM. We validated our surrogate embeddings on two qualitatively different downstream tasks: protein-protein interaction (a pairwise binary task) and subcellular localization (a single-protein ten-class task). In both cases, surrogate embeddings demonstrate minimal degradation in downstream task performance with a significant reduction in memory footprint. In future work, we want to extend this work to other PLMs beyond ESM and explore piecewise polynomial surrogates for improved reconstruction. We also aim to extend the surrogate framework to per-residue embeddings and investigate if whether the smooth-trajectory assumption holds for per-residue embeddings, enabling residue-level tasks such as secondary structure prediction, and to evaluate on additional protein-level tasks such as multi-label localization.

## Acknowledgement

We would like to acknowledge computing resources provided by the Center for Advanced Research Computing (CARC) at the University of New Mexico.

## A Appendix

To validate our hypothesis that protein-level (mean-pooled) PLM embeddings follow a predictable path across different transformer layers. We examine their behavior through first two PCA components corresponding to each layer for four representative proteins. Figure 4 illustrates the first two PCA components of each layer for both real(solid lines) and reconstructed embedding(dashed lines). The selected proteins represent different levels of overall reconstruction quality: (a) Q96GC6 at the 90th percentile, (b) Q96N77 at the median, (c) Q7Z5J4 at the 25th percentile, and (d) O14904 at the 10th percentile.

**Figure 4:**
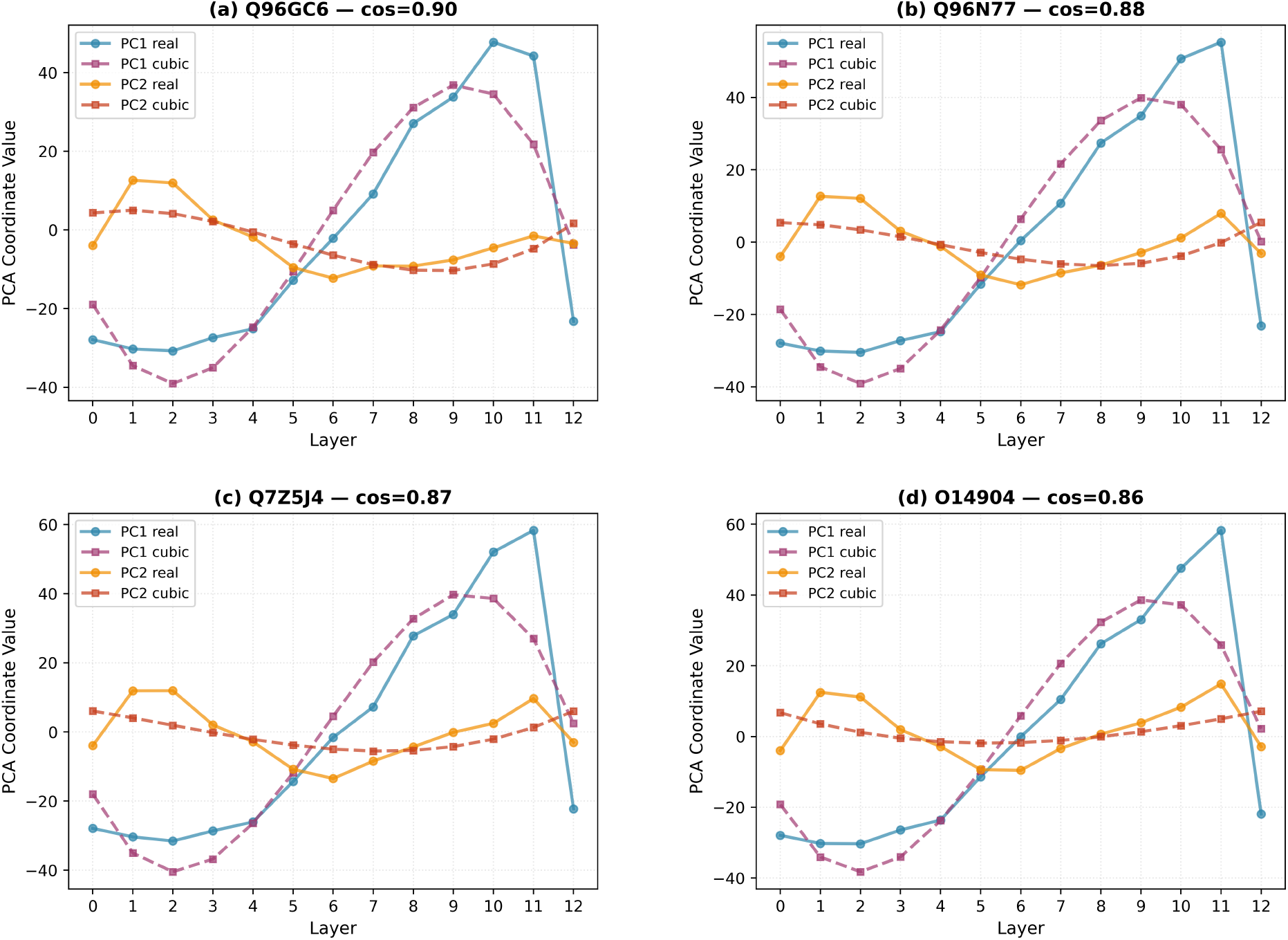
First two PCA components across layers for real (solid lines) and reconstructed (dashed lines) embeddings for proteins corresponding to different reconstruction quality.

Figure 5 depicts the performance at layer 7 for the ESM-35M PPI task. This demonstrates the robustness of the task across multiple layers.

**Figure 5:**
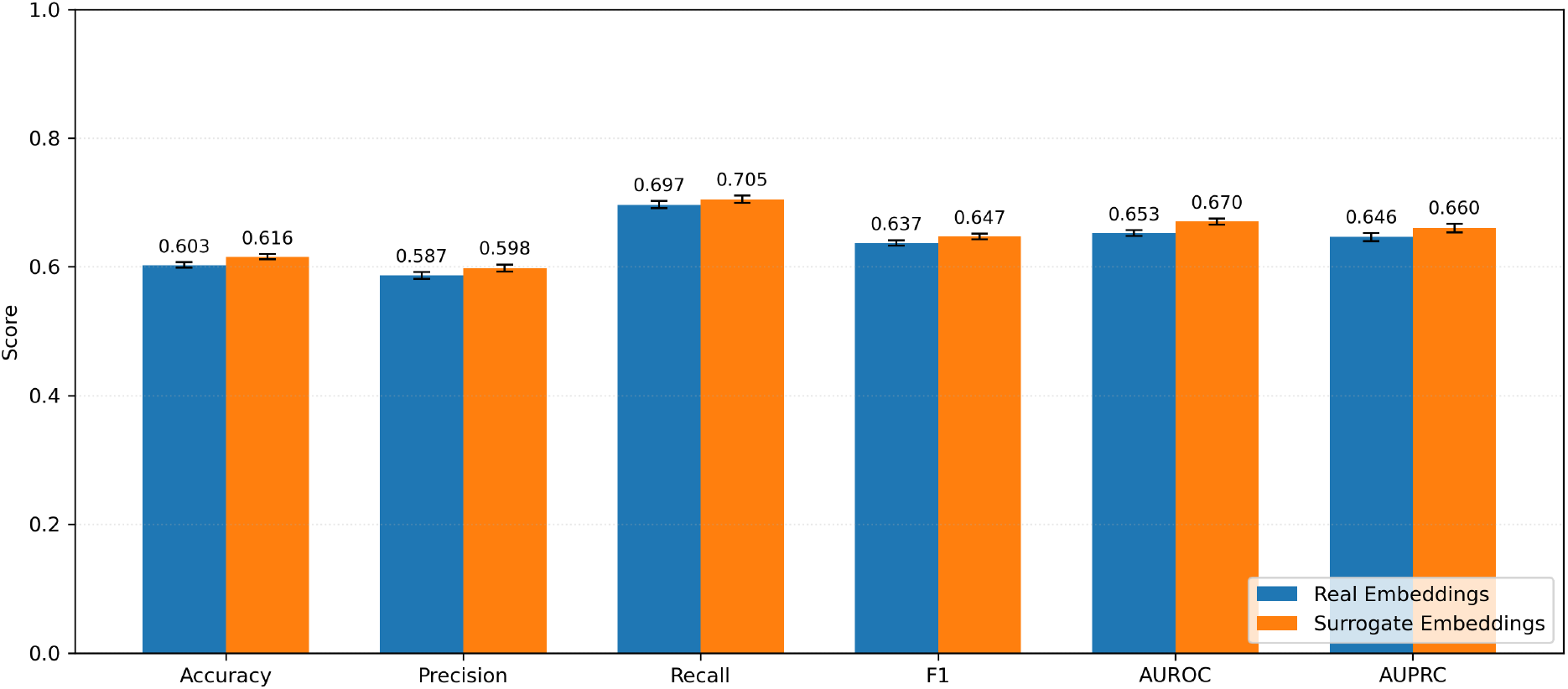
Test-set performance of original and surrogate embeddings for PPI prediction task at layer 7. Error bars denote 95% bootstrap confidence intervals (1000 resamples).

Figure 6 shows the layer selection for ESM-3B model using the validation dataset. The best selected layer based on our metrics (AUROC) is layer 32, marked by the dashed line. The final layers show some drop in performance that further supports our motivation that the last transformer layer is not always optimal for all downstream tasks

**Figure 6:**
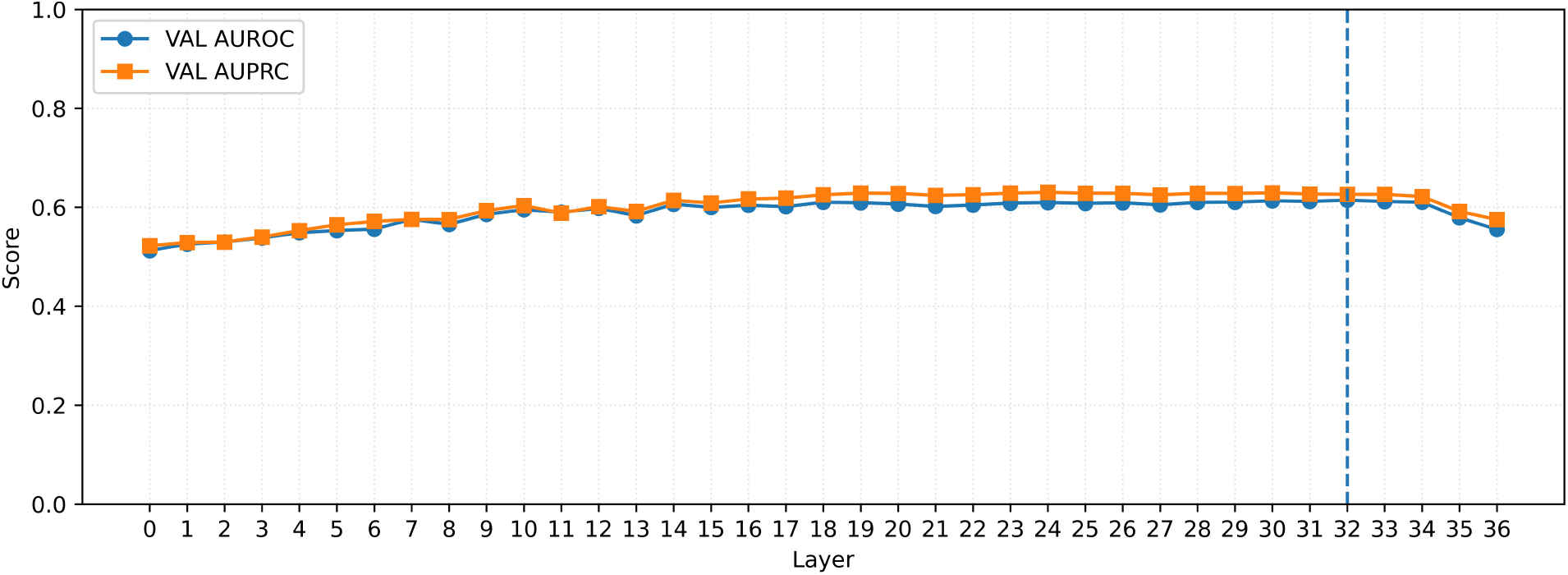
Layer Wise PPI prediction performance on validation data using ESM-3B.

### A.1 Comparison to standard compression baseline

We also compared our surrogate against two baselines on both downstream tasks: FP16 quantization and PCA only projection at the same *K* used by our surrogate method. All methods perform comparably across both tasks (AUROC within 0.02 compared to full FP32 embedding), confirming that PLM embeddings are highly compressible. The key difference is with storage (compressing ratios): FP16 always yields 2*×* compression. PCA-only at *K* dimensions yields a *D/K* ratio: 3.7*×* for subcellular localization on ESM-35M (*K*=128, *D*=480), 7.5*×* for PPI on ESM-35M (*K*=64), 20*×* and 40*×* on ESM-3B (*D*=2560). Our surrogate adds one more compression step. Rather than storing the coordinates at every layer, we fit a polynomial across layers and store just its *p* coefficients (*p* = 4): 12*×* for subcellular localization with ESM-35M, 24*×* for PPI ESM-35M, 144*×* for subcellular localization with ESM-3B, and 360*×* for PPI ESM-3B. This gets us to 12 − 144*×* overall compression.

